# Long-TUC-seq is a robust method for quantification of metabolically labeled full-length isoforms

**DOI:** 10.1101/2020.05.01.073296

**Authors:** Sorena Rahmanian, Gabriela Balderrama-Gutierrez, Dana Wyman, Cassandra Joan McGill, Kim Nguyen, Robert Spitale, Ali Mortazavi

**Affiliations:** Department of Developmental and Cell Biology, University of California Irvine, Irvine, CA. 92697, USA; Center for Complex Biological Systems, University of California Irvine, Irvine, CA 92697, USA; University of California, Irvine, Department of Pharmaceutical Sciences, Irvine, CA 92697, USA

## Abstract

The steady state expression of each gene is the result of a dynamic transcription and degradation of that gene. While regular RNA-seq methods only measure steady state expression levels, RNA-seq of metabolically labeled RNA identifies transcripts that were transcribed during the window of metabolic labeling. Whereas short-read RNA sequencing can identify metabolically labeled RNA at the gene level, long-read sequencing provides much better resolution of isoform-level transcription. Here we combine thiouridine-to-cytosine conversion (TUC) with PacBio long-read sequencing to study the dynamics of mRNA transcription in the GM12878 cell line. We show that using long-TUC-seq, we can detect metabolically labeled mRNA of distinct isoforms more reliably than using short reads. Long-TUC-seq holds the promise of capturing isoform dynamics robustly and without the need for enrichment.

## INTRODUCTION

Transcription is a dynamic process and different transcriptome profiles are indicative of different cellular states. While each cellular state can be identified by a set of quasi-steady state expression levels, all mRNA transcripts are transcribed and degraded at different rates ^1,2^. The expression level of each gene isoform depends on its transcription rate, processing rate, and degradation rate. Although regular RNA-seq studies inform us of the steady state levels of each transcript, these lack any information on transcript stability or turnover rates. Transcription is controlled by cis-regulatory elements such as promoter and enhancer regions which play a role in determining the transcription rate of a transcript ^3^. The binding of transcription factors as well as characterization of epigenetic marks from this category is primarily studied using ChIP-seq ^4^ and the chromatin accessibility can be measured by assays such as ATAC-seq ^5^. However, RNA degradation rates are just as important, and often times overlooked, when defining the steady state levels of expression (Maekawa et al.; Ghosh and Jacobson). Post-transcriptional regulatory factors such as miRNA and RNA binding proteins are the main players in regulating RNA stability and decay. Assays such as CLIP-seq and miRNA-seq have been developed to study the effects of each of these elements on gene expression ^8,9^. Overall, transcription is a complex process and using the expression profiles to understand the role of each of these regulators can be ambiguous and challenging.

Several new methods have been developed for genome-wide study of transcription dynamics. One category of these methods focuses on the study of nascent transcriptomes by profiling the RNA molecules instantaneously as they are being transcribed or processed. For instance, global run-on sequencing (GRO-seq) and precision nuclear run-on sequencing (PRO-seq) sequence the positions that the polymerase is residing at, providing information regarding active genes and the polymerase pausing dynamics ^10,11^. Another set of methods, such as native elongating transcript sequencing (NET-seq), report polymerase positions at the 3’ ends of nascent transcripts ^12,13^. While GRO-seq, PRO-seq, and NET-seq investigate nascent transcripts, other methods focus on metabolic labeling of nascent RNA molecules that have been made over a window of time in order to study transcription and degradation rates. These methods use different nucleotide analogs to label the newly made RNA over a pulsing window followed by high throughput sequencing to detect the RNA molecules that incorporated the analog. A group of these methods such as bromouridine sequencing (Bru-seq), 4-thiouridine sequencing (4SU-seq) and transient transcriptome sequencing (TT-seq) rely on enrichment methods to recover signal from labeled transcripts ^14–16^. Many of these methods suffer from enrichment biases and elution issues that lead to low yield and biases due to modified nucleotide identity used for enrichment.

More recently, additional methods have been developed that can still characterize modified nucleoside incorporation, but do not rely on enrichment. TimeLapse-seq, thiol(SH)-linked alkylation for the metabolic sequencing (SLAM seq), and thiouridine to cytidine conversion sequencing (TUC-seq) rely on chemical conversion of the metabolically incorporated analog. Modified positions are then identified in mutated cDNA in order to distinguish the metabolically labeled reads from pre-existing none-labeled reads ^17–19^. One of the challenges of this group of methods is the low incorporation rate of 4SU that results in under-estimation of recently transcribed genes ^20^, especially when using short-read sequencing, which is still a long-standing challenge in transcriptomics, especially when interrogating more complex transcriptomes with large dynamic range.

All of these techniques rely on short-read Illumina sequencing, which even with higher sequencing depth cannot overcome these limitations. In addition, reconstructing different transcript models and quantifying the expression at the level of isoforms using short reads remains challenging and limited ^21^. Long-read sequencing can improve the sensitivity of the assay by sequencing over the whole transcript, which would have a higher number of 4SU incorporated and makes it easier to detect over sequencing and biological noise. The two main long-read sequencing platforms are Pacific Biosciences (PacBio) and Oxford Nanopore Technology (ONT). Despite the higher error rates in long-read technologies, the circular consensus technique implemented by PacBio has reduced the final error rate down to 1% ^22^. Furthermore, long-read sequencing can unambiguously identify transcript isoforms using packages such as TALON (Wyman and Balderrama-Gutierrez et al., 2019), SQANTI ^24^ or FLAIR ^25^.

In this work, we combine TUC metabolic labeling with long-read sequencing on the PacBio Sequel II platform to develop long-TUC-seq. We pulsed the GM12878 cells with 4 thio-Uridine (4SU) for 8 hours and then converted the incorporated 4SUs into cytidines using osmium tetroxide. We then built cDNA and libraries for sequencing on both Illumina NextSeq and PacBio platforms. We quantified the expression levels of each gene that correspond to the recently made RNA during the 8 hours pulsing window by quantifying the number of T→C substitutions identified in every read. We explored different thresholds to count the read with different levels of certainty as newly synthesized. We demonstrate that long-TUC-seq has higher sensitivity and lower FDR compared to the corresponding short-read version of TUC-seq. Finally, we count the reads in each category for all the isoforms to identify differences in transcription rates between isoforms of the same gene. Overall, long-TUC-seq is a robust protocol that would be widely applicable to a variety of settings were the metabolic labeling can be used to study transcriptome dynamics.

## METHODS

### Sample collection and RNA extraction

GM12878 cells were obtained from Corriell Institute and were cultured in accordance with ENCODE protocols (www.encodeproject.org). The cells were passed every two to three days at 200k-500k cells/mL density and were harvested for the experiments at 500k-1M cells/mL. The RNA was extracted using QIAGEN RNeasy Plus kit (Cat. No. 74134).

### TUC-seq sample preparation

4-thiouridine was obtained from Sigma Aldrich (T4609) and used fresh at a working concentration of 200 mM. For each TUC-seq experiment, 10-15M cells were spun down and resuspended in 10-15 mL of fresh media with added 4SU at a final concentration of 1mM (no 4SU was added for the osmium controls). The cells were incubated with 4SU for 8 hours and harvested for RNA extraction. The RNA was then treated with OsO_4_ solution for 3 hours at room temperature in dark. The osmium solution was prepared fresh every time by mixing 20 μl of 1mM OsO_4_ (Sigma Aldrich, 201030) with 4μl of 2M NH_4_Cl at pH 8.8 and 1μl of RNasin Plus RNase inhibitor (Promega, N2615) for every 10μg of RNA. The RNA was then purified using Zymo RNA cleanup kit (R1015). Finally, the RNA was treated with 1U of exonuclease from epicenter (Terminator™ 5’-Phosphate-Dependent Exonuclease, TER51020) for 1 hour at 30°C and neutralized by 1μl 100mM EDTA. Then, the RNA was once more purified with Zymo RNA cleanup kit.

### PacBio library preparation and sequencing

The set III of SIRV controls were spiked into the RNA samples at a level of 0.03% of the total RNA. The cDNA was generated using a modified version of SMART-seq2 protocol. We then followed SMRTbell Template Prep Kit 2.0 to build PacBio libraries using 1-2μg of input RNA. We checked the quality of the libraries using the Bioanalyzer 2000 and Qubit to get the final concentrations. Finally, the libraries were delivered for sequencing on a Sequel II platform at UCI sequencing core facility, using 1 SMRT cell per library.

### Illumina library preparation and sequencing

Starting from 30-50ng of the same cDNA, we followed the Illumina tagmentation protocol using Nextera DNA Flex Library Prep Kit to generate Illumina short-read libraries. We checked the concentration of the libraries with Qubit and got the average length of the library using the Bioanalyzer. We then performed a 2×43 paired-end sequencing on our NextSeq 500 instrument.

### PacBio data processing

Raw reads from Sequel II machine were processed by PacBio circular consensus package (CCS v4.0.0) to filter any reads with less than 3 passes (parameters: --noPolish --minLength=10 -- minPasses=3 –min-rq=0.9 –min-snr=2.5). Then reads with misconfigured adapters were filtered using PacBio lima package (v1.10.0; parameters: --isoseq --num-threads 12 --min-score 0 --min-end-score 0 --min-signal-increase 10 --min-score-lead 0). Finally, full-length non-chimeric (FLNC) reads were extracted using the PacBio Refine package (v3.2.2; parameters: --min-polya-length 20 --require-polya). The bam files processed by Refine were then converted to fastq files and they were all deposited to GEO (https://www.ncbi.nlm.nih.gov/geo/query/acc.cgi?acc=GSE149551) with the exception of PacBio GM12878 control sample which has been previously deposited onto ENCODE portal (https://www.encodeproject.org/experiments/ENCSR838WFC/).

The FLNC reads were then aligned to a modified version of human genome reference (GRCh38 with added SIRV and ERCC references) using minimap2 (v2.17; parameters: -ax splice:hq -t 16 --cs -uf). We then used TransciptClean (v2.0.2; parameters: -m False --primaryOnly) for reference-based error correction of the reads. We provided TranscriptClean with splice junctions reference derived from the GENCODE annotations using TranscriptClean accessory script get_SJs_from_gtf.py. We also provided it with VCF-formatted NA12878 truth-set small variants from Illumina Platinum Genomes. We first initialized the TALON database with GENCODE v29 + SIRVs/ ERCC annotations using talon_initialize_database and finally annotated the reads by running TALON V4.4.2 module on all the datasets. We obtained the table of annotated reads from all the datasets by running the talon_summarize module. All the scripts used for analysis of long-TUC-seq Pacbio data can be accessed on the mortazabilab github. (https://github.com/mortazavilab/long-TUC-seq)

### PacBio labeling of the reads

We used a custom python script (mismatch_analysis_PB.py) to annotate the reads with their corresponding substitutions. The script uses the CS tag option from minimap2 to count different types of substitutions and to generate a text file containing each read name and its corresponding substitution tally. The script also breaks down the alignment file into subfiles, each containing one of one category of reads (> 0, > 6, > 20 and > 30 T→C) for visualization on the UCSC genome browser. The information on different substitutions was added to the annotations obtained from TALON. We then calculate the TPM and counts for each of the categories for each gene and transcript.

### Illumina data processing

The reads from illumina runs were mapped to human transcriptome reference (GRCh38.p12, gencode.v29.primary_assembly.annotation) using STAR aligner (v2.6.0c; parameters: -- outFilterMismatchNmax 15 --outFilterMismatchNoverReadLmax 0.07 -- outFilterMultimapNmax 10 --outSAMunmapped None --outSAMattributes MD NM --alignIntronMax 10 --alignIntronMin 20 --outSAMtype BAM SortedByCoordinate). The raw fastq files for each sample is available on GEO database under GSE149551 accession. (https://www.ncbi.nlm.nih.gov/geo/query/acc.cgi?acc=GSE149551).

### Illumina calling of the labeled reads

We ran a custom python script (mismatch_analysis_ill.py) to annotate each of the mapped reads with the number different substitution events. The script uses the MD tag to tally the number of substitutions for each read. The script also breaks down the alignment file into sub-files of reads with > 0, > 2, > 4 and > 6 T→C substitutions. Finally, we count the reads in each category using eXpress (v1.5.1; parameters: --no-bias-correct). The quantification can be accessed under GSE149551 accession in GEO database. (https://www.ncbi.nlm.nih.gov/geo/query/acc.cgi?acc=GSE149551). All the scripts used to process illumina TUC-seq data can be accessed on the Mortazavilab github. (https://github.com/mortazavilab/TUC-seq)

### Degradation rate and half-life calculations

Assuming steady-state and doubling rate of zero during the pulsing time, we can calculate the degradation rate (λ_i_) and consequently the half-life (hl_i_) of gene i:

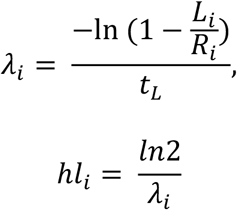

Here R refers to the steady state expression of the specific mRNA, L stands for the expression of labeled RNA, and t_L_ is the labeling time.

### Isoform specificity analysis

In order to help us understand the isoform specificity of each gene and its dynamics, we introduce an index for isoform specificity of a gene (ISI) as follows:

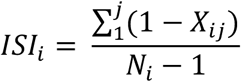

Here, index *i* corresponds to each gene and index *j* represents each corresponding isoform for gene *i*. X is the expression level of the isoform normalized to the expression level of the highest expressed isoform of the gene *i*. Finally, N_*i*_ is the number of isoforms corresponding to gene *i*. We calculated the isoform specificity indices for each of the genes using the total and labeled RNA. Then we filter for the genes with more than 2 isoforms that has an ISI_total_ < 0.35 and ISI_labeled_ > 0.85. We then plot the expression of each isoform of a representative set of these genes and color the portion of the expression that corresponds to the labeled reads.

## RESULTS

### Identifying metabolically labeled RNA using long-TUC-seq

Our long-TUC-seq method relies on the incorporation of 4SU into the RNA and its further conversion to a regular cytidine (Fig. 1A.) We initially tested 4SU incorporation into recently synthesized transcripts by incubating GM12878 cells with 0.1mM and 1mM 4SU for a period of time between 2 to 24 hours and compared the amount of incorporation by dot blots. We then checked the RNA integrity after the treatment of the RNA samples with osmium tetroxide under different conditions (mainly time and temperature of the incubation). We compared the RNA Integrity Numbers (RINs) of the RNA samples after the treatment using a Bioanalyzer. Even with milder temperature (room temperature) we observed substantial degradation at 3 hours (RIN = 5.6). However, the integrity of the samples is improved with the addition of RNase inhibitor to the OsO_4_ mix at this condition. Finally, we tested the conversion of incorporated 4SU by OsO_4_ at this condition by checking the amount of 4SU remaining in the RNA sample before and after osmium treatment, using a dot blot assay. The dot blot shows complete conversion of 4SU with 3 hours of OsO_4_ at room temperature.

**Figure 1.**
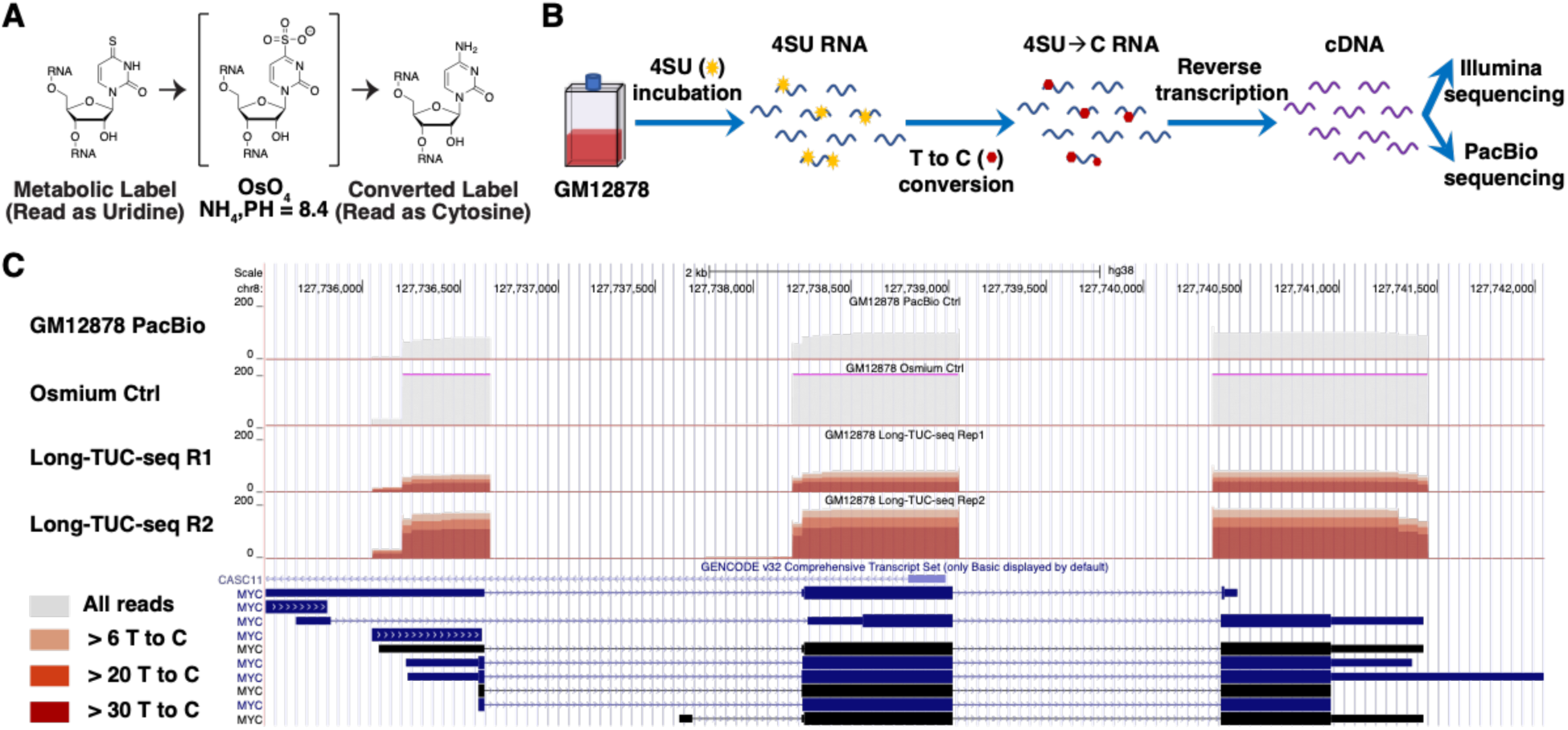
Identification of recently synthesized transcripts in GM12878 by long-TUC-seq. Osmium tetroxide converts an incorporated 4SU into a regular cytidine. **b)** Experimental layout of TUC-seq sample preparation, starting with the incorporation of 4SU into the GM12878 cells following by its conversion to C using OsO_4_ and finally library building from cDNAs. **c)** Genome browser screenshot of PacBio data of GM12878 control from ENCODE, Osmium treated GM12878 without 4SU incorporation and two biological replicates of long-TUC-seq samples. The shot shows reads aligned to MYC, with increasing levels of labeled reads colored with darkening shades of red. The tracks are shown on a scale of 0 to 200 reads.

We pulsed biological replicates of GM12878 cells with 1mM of 4SU for 8 hours and extracted the RNA, which were treated with osmium tetroxide. We also generated matching libraries of osmium treated samples without any 4SU pulsing. We built Illumina and PacBio libraries from these samples and sequenced them on their respective platforms and analyzed the data (Fig. 1B). Each of the PacBio libraries yielded between 3.4M - 6.2M raw sequencing reads (Table S1). After all the filtering, we are left with a minimum of 1.2M of reads for each sample that were mapped to human genome using minimap2 with an average of 99.65% mapping rate. In order to identify the reads that were synthesized during the 4SU pulse window, we counted the number of T→C substitutions for each read. We inspected the reads that mapped onto the MYC locus, which is known to be a fast turnover transcript (Fig. 1C). We observe that a high percentage (94%) of TUC-seq reads mapping to the MYC locus have at least 6 T→C events. By contrast, none of the reads mapping to MYC in the osmium control (sample without 4SU pulse and treated with OsO_4_) or in publicly available PacBio ENCODE datasets are marked as labeled. We can therefore detect 4SU labeled reads based on the number of substitutions in a long read.

### Distinct substitution profiles of long-TUC-seq at the level of base calls and reads

The nucleotide composition of the human genome is equally distributed between all the four nucleotides. There are some biological variations from multitude of SNPs that will introduce specific substitution events across the genome and some technical variation that is introduced via PCR or SBS. However, all of these substitutions should be distributed evenly between the 12 different possible substitution types globally. While this equal distribution is observed in the control PacBio RNA-seq from ENCODE and the osmium control, there is a very distinct profile in our long-TUC-seq samples with a much higher T→C counts as expected (Fig. 3A). In order to asses our ability to call a read as labeled, we analyze the distribution of reads based on the number of T→C observed. We detect 34% of all the reads being labeled with more than 6 T→C in the TUC-seq samples compared to 0.4% in the osmium control and in the RNA-seq control (Fig. 3B). In addition, we detect 27% and 21% of the reads from the TUC-seq samples are labeled with a minimum of 20 and 30 T→C.

**Figure 2.**
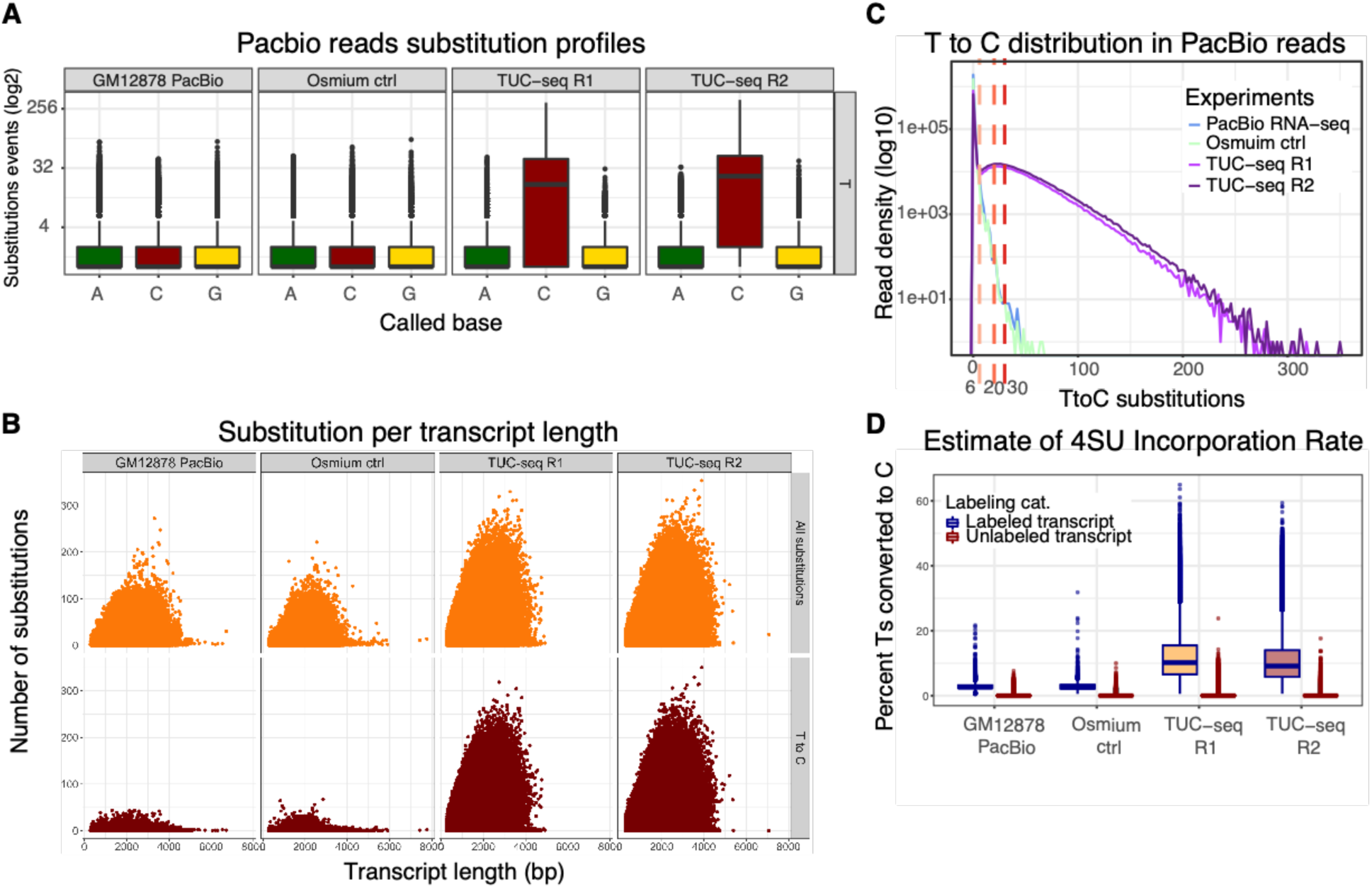
Identifying labeled reads in long-TUC-seq. **a)** Comparison of the profiles for all the possible substitutions at a reference T base across the TUC-seq samples and controls. **b)** The distribution of PacBio reads with respect to the number of T→C substitutions observed for each read. The three dotted lines indicate the thresholds we used to define lower, medium and higher labeled reads. **c)** Number of substitution events observed in each read with respect to the length of each read, showing slightly higher T→C events in the longer reads for TUC-seq samples. **d)** Average number of Ts in each read converted to C for labeled and unlabeled reads across the TUC-seq samples and controls.

**Figure 3.**
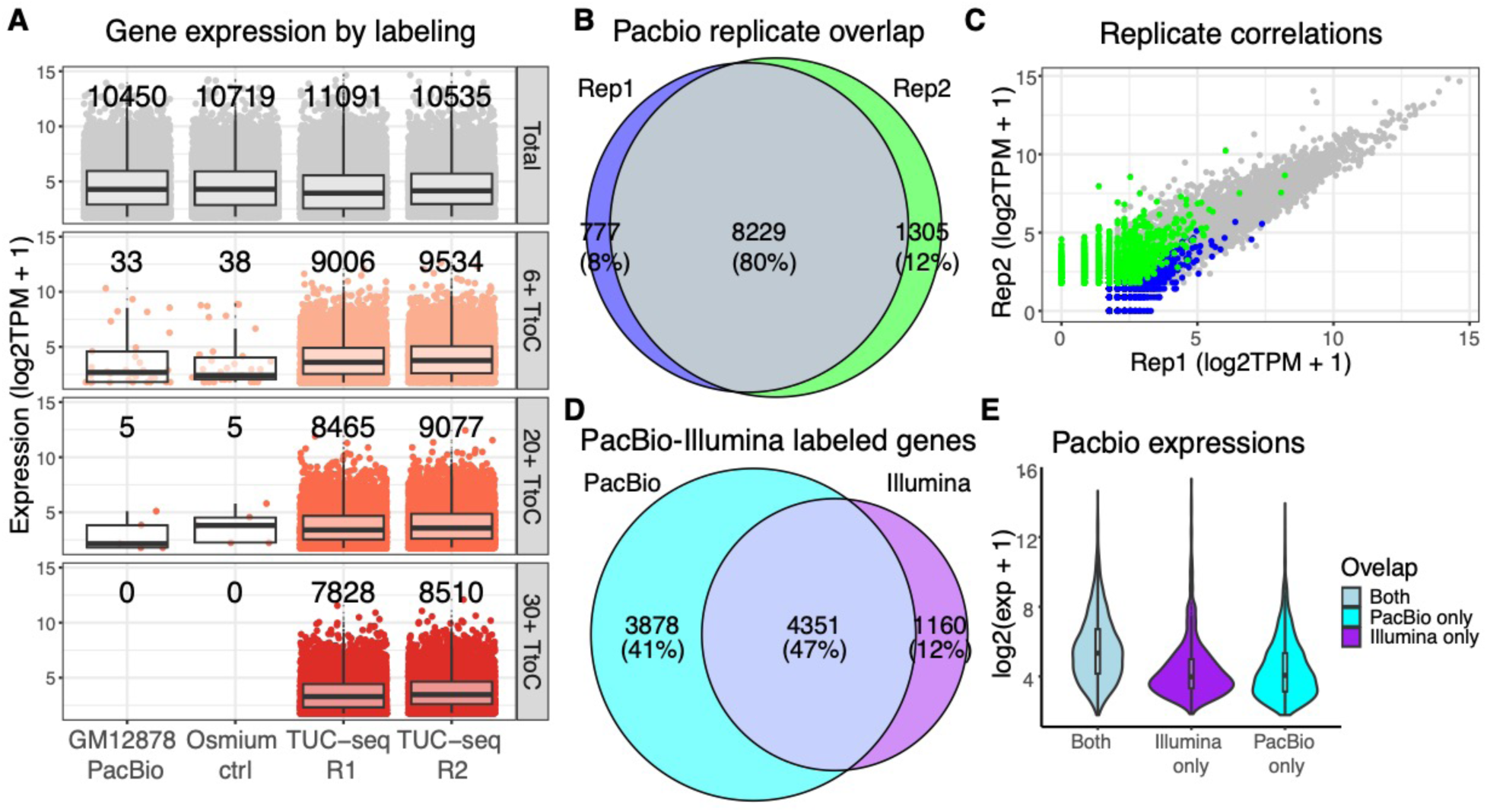
Robust identification and quantification of recently transcribed genes. **a)** Expression levels of genes with more than 2 TPM in each of the labeling categories. The total number of genes is indicated on top of each of the boxplots. **b)** The overlap of genes detected as lower labeled in either of the TUC-seq replicates showing the percentage of genes in each section of the Euler diagram. **c)** The correlation of lower labeled genes between the two replicates with a Pearson correlation coefficient of 0.93. **d)** Overlap of genes detected as lower labeled by both replicates on Illumina and those detected by both replicates of PacBio. **e)** The expression levels of the genes in each section of the Euler plot in section c. The expression levels of “both” and “PacBio only” groups are from PacBio and the expression levels of the middle group (“Illumina only”) is from Illumina data.

To ensure that the reads labeled by long-TUC-seq are not heavily biased by longer transcripts, we determined the correlation of the number of T→C with the length of each transcript. Although the number of observed T→ C in a read does correlate weakly with length of the transcript (Pearson correlation coefficient of 0.25), its distribution in the controls indicates that the transcript length is not a big driver of noise, which will therefore not hinder an accurate count of labeled transcripts (Fig. 3C). Finally, we counted the number of Ts in each transcript that has been converted to C in order to obtain an estimate of 4SU incorporation rate. Our 8-hour long-TUC-seq results indicate an average of 11.33% for 4SU incorporation in the transcription process, assuming a 100% conversion to C (Fig. 3D).

### Robust detection of recently synthesized genes by long-TUC-seq

We used TranscriptClean ^26^ to correct the indels in our reads before running TALON V4.4.2 (Wyman and Balderrama-Gutierrez et al., 2019) to annotate the reads as known and novel transcripts, as well as to obtain accurate counts for each gene and transcript for each of our 4 datasets (1 RNA-seq control, 1 osmium controls and 2 TUC-seq samples). For the purpose of this study, we focused on known isoforms. We detect 21,496 known genes across the experiments and 32,250 (TPM > 0) known transcripts. We added the labeling information for each read to the TALON annotations and calculated the expression levels for each gene and transcript for the following 4 categories: all reads, permissive threshold (>6 T→C), intermediate threshold (>20 T→C) and conservative threshold (>30 T→C.) We detect an average of 9,270 genes labeled at permissive threshold with more than 2 TPM expression of labeled reads, in the TUC-seq samples compared to 35 genes out of 10,584 genes detected in the controls (FDR = 0.33%). This number drops to 8,169 in the conservative category of labeled reads in the TUC-seq samples (Fig. 3A). The detection of recently synthesized genes is very robust across the replicates, with 80% of detected labeled genes (> 2 TPM at permissive threshold) being confirmed by both replicates (Fig. 3B). There is also a high concordance amongst the expression levels of these recently synthesized genes across the replicates with 0.93 Pearson correlation (Fig. 3C). This correlation is still high for genes detected in the higher categories with Pearson correlation of 0.93 for intermediate labeled reads and 0. 92 for conservative reads.

### Comparison of long-TUC-seq with Illumina short-TUC-seq

Current methods using metabolic labeling for studying the dynamics of transcription rely on short read illumina sequencing. In order to benchmark our long-TUC-seq results we compared it with the short-read TUC-seq of the same samples. We built the Illumina Nextera libraries using the same cDNA materials that were used for PacBio libraries. We then sequenced these libraries on the Illumina NextSeq platform and mapped the reads to the human transcriptome reference using STAR with an average of 45M single end reads mapped per sample. We annotated each read with the number of observed substitutions and annotated the aligned reads with it. Here we also detect higher T→C substitution profile for TUC-seq samples compared to the controls. The TUC-seq samples contain more reads with higher T→C compared to the controls; based on the substitution profiles and the read distributions, we decided to used 2, 4 and 6 T→C as permissive, intermediate and conservative thresholds for calling the labeled reads. We detect 27% of total reads labeled with > 2 T→C in TUC-seq samples compared to 1.5 % in control samples. Although raising the threshold to 4 T→C reduces the percentage of false positive labeled reads in controls to 0.14%, it also reduces the percentage of labeled reads in the TUC-seq samples to 15%. Finally, we calculated the 4SU incorporation rate from Illumina short-read TUC-seq samples to be 17.22% which is 6% higher than what we have obtained using Pacbio long-TUC-seq data.

Using the permissive threshold of 2 T→C, we detect 57% of reads mapping to MYC in labeled samples, which is 37% lower than what was detected by PacBio. We then quantify the expression levels in each category using eXpress ^27^ as described in the methods. In order to compare the detection of labeled genes by each platform, we use the intermediate threshold for Illumina (4 T→C) which resulted in similar FDR (0.5%) to that of PacBio data with permissive threshold (FDR = 0.3%). Although Illumina TUC-seq detects twice as many genes across all the samples compared to PacBio (> 0 TPM), the number of detected genes at intermediate threshold is 5,511, which is much less than labeled genes in PacBio. When comparing genes (expressed > 2 TPM) detected as labeled in either platforms, we find that 47% are shared and the majority of the remainder (41% of all labeled genes) is detected only in PacBio (Fig. 3D). In general, the expression levels of the genes detected at 2 TPM or higher by only one of the platforms is lower than the expression of the commonly shared detected genes (Fig. 3E). Thus long-TUC-seq is more sensitive than its short-read equivalent at similar FDR thresholds.

### Calculating degradation rates with long-TUC-seq

After annotating the detected genes with the different degrees of labeling, we focused on the dynamics of transcription for each gene and analyzed the rate at which each gene is transcribed. On average, 50% of the total expression of genes at the end of our 8-hour labeling window comes from newly synthesized RNA. MYC is one of the genes with faster turnover rate that is expressed at 111 TPM with 95% of its expression being labeled whereas GAPDH with a high expression of 11,378 TPM has only 5.6% labeled RNA (Fig. 4A; Table S2).

**Figure 4.**
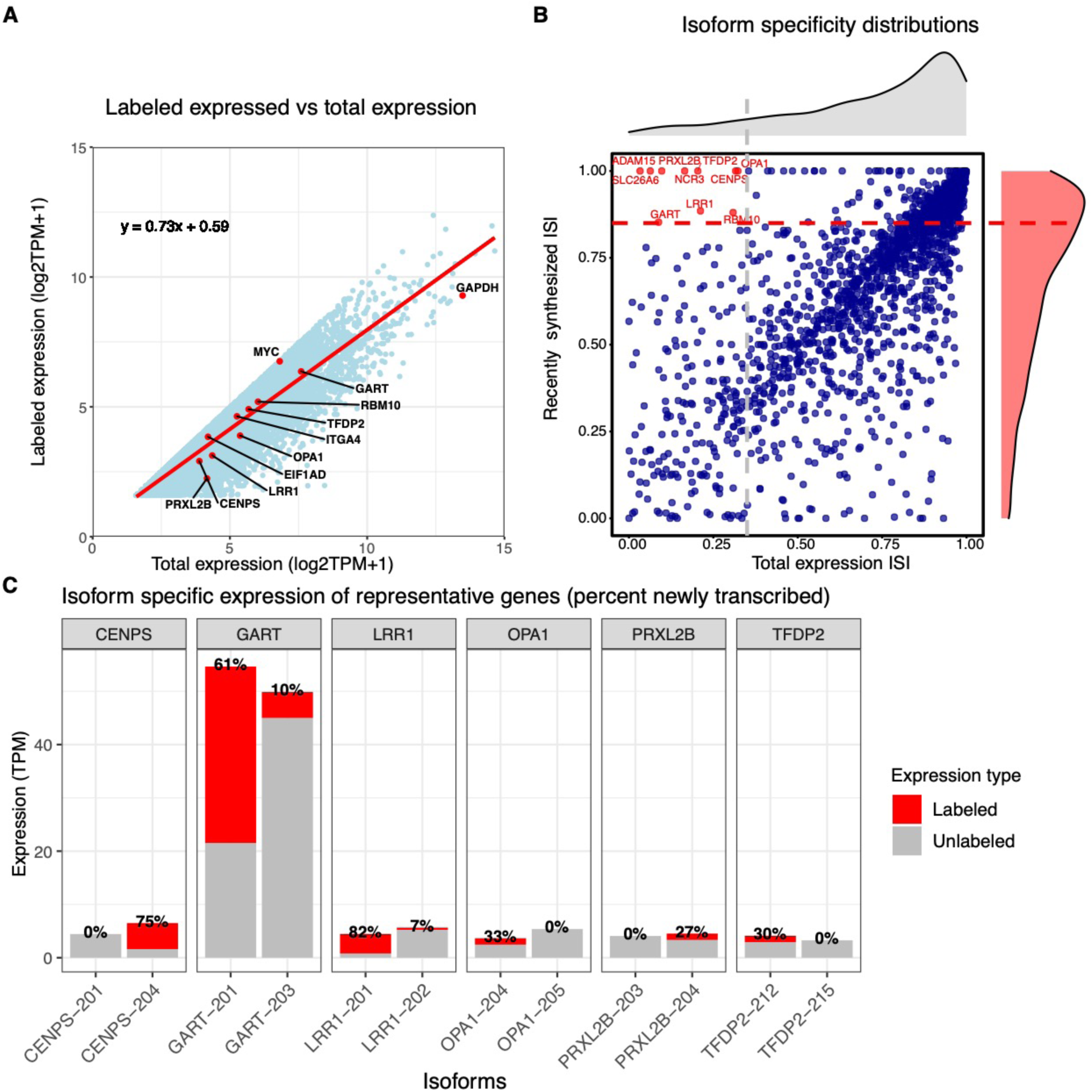
Dynamics of expression at the level of individual isoforms. **a)** Expression levels of recently transcribed genes (labeled at permissive threshold) with respect to the total expression level of that gene (for genes >2 TPM). The equation corresponds to the regression line drawn in red. Two example genes (MYC and GAPDH) are highlighted in red. **b)** The distribution of isoform specificity indices for all of the genes calculated from total expression (in grey) and from recently made transcripts (in red). The dotted lines indicate the thresholds used to find genes with lower ISI_total_ (< 0.35) and higher ISI_labeled_ (> 0.85). **c)** Expression levels of the isoforms corresponding to representative genes from the set defined in b. The grey portion of the bars corresponds to the expression level of pre-existing RNA and the red portion corresponds to the recently synthesized transcripts. Finally, the percentages on top of the bars are representing the percentage of total expression of the isoform that is transcribed recently.

Under a steady-state assumption that the overall expression level of a gene stays the same through the 8-hour pulsing window, the rates at which a gene is being transcribed and the rate at which it is degraded are constant. We calculated the degradation rates and the half-life of each gene using the total expression of the gene and its newly synthesized RNA. We obtained a degradation rate of 45 TPM/hour and a half-life of 1.7 hours for the MYC gene. The ranking of genes based on their half-life time is similar to what has been observed previously (Spearman correlation of 0.74 with timeLapse-seq ranking in K562 cells) ^17^. Although we used a long labeling time of 8 hours, the method could work with substantially shorter labeling time. Long-TUC-seq can be used to calculate degradation rates from 4SU labeling of transcripts and genes.

### Analysis of isoform-specific expression and transcription rates

One of the advantages of long-read sequencing is that it inherently measures the expression levels of the isoforms of each gene. In our study, more than 58% of genes are expressed as multiple isoforms with an average of 2.5 isoforms per gene. GAPDH, which is one of the higher expressed genes, has 4 distinct isoforms. MYC, which is one of the higher turnover genes, has 2 isoforms detected. The highest number of isoforms belong to MSL3, with 15 isoforms detected. We can also take a step further and analyze the expression levels of each isoform to see if the gene is expressed through one isoform more than the other, or if it is expressed uniformly across different isoforms by calculating the isoform specificity index (ISI) for all genes as described in the methods. In the case of a gene that expresses all its isoforms equally, the ISI will be closer to zero and in the case of a gene that expresses primarily one of its multiple isoforms, the ISI will be closer to one. MYC and GAPDH each have an ISI of 0.67 and 0.99, respectively, which for MYC translates to the fact that its isoforms are expressed in a 3:1 ratio, and for GAPDH it means that its isoforms are expressed in approximately a 800:20:4:1 ratio.

We can similarly define the isoform specificity index based on the expression levels of newly synthesized transcripts (ISI_new_) and inspect the isoform specificity of the transcription machinery for a specific gene at a given time. The distribution of ISI_total_ and ISI_new_ for all the genes of GM12878 shows that majority of multi-isoform genes are expressed and being transcribed in a highly isoform-specific manner, and there is a Pearson correlation coefficient of 0.63 between total and labeled isoform specificity (Fig. 4B; Table S3). Furthermore, we are interested in genes with ubiquitous isoform expression that are being transcribed in an isoform-specific manner. In order to obtain a list of such genes, we filter the genes with lower ISI_total_ (< 0.35) and higher ISI_new_ (>0.85). There are 9 genes in this category, all with two isoforms detected in our dataset, the expression of which are less than two-fold apart. However, the expression of recently synthesized isoforms is in some cases more than 70-fold different (Fig. 4C). One such gene is LRR1 that encodes for Leucine-rich repeat protein 1, which plays a role in protein ubiquitination and modification. This gene has five isoforms, two of which have been detected in our dataset with similar expression levels of about 5 TPM (201 and 202). These isoforms are protein coding and they differ only in one exon; however, the LRR1-202 isoform which has an extra exon compared to the 201 isoform has a much higher turnover, and about 73% of its expression has been made within the 8-hour pulse window.

## DISCUSSION

Here, we introduce a method for detecting and quantifying metabolically labeled RNA at a single isoform resolution using PacBio long-read sequencing. To do so, we relied on the conversion of incorporated 4SU to C by TUC-seq chemistry. We demonstrated that even though short-read Illumina sequencing provides much higher depth in comparison to PacBio sequencing, we are able to recover higher number of labeled genes with PacBio. We also show that not only can PacBio detect the labeled RNA reproducibly, the quantification of these labeled RNAs is also highly concordant between the biological replicates. Furthermore, we took advantage of having T to C substitution data for full transcripts in order to calculate an accurate estimation of 4SU incorporation rate within each transcript. This estimation using illumina short-read technique would be in accurate and over-estimated due to the fact that many of the reads aligning to the T depleted regions are dis-missed as unlabeled.

We use long-TUC-seq data to obtain estimations of degradation rates of genes and consequently their half-lives. The caveats with these estimations are the two assumptions used in their calculations. First is the steady state assumption that the expression level, synthesis rate, and degradation rate of each gene is constant during the pulsing time, which can be closer to reality when the pulsing time is much shorter than 8 hours. The other assumption used in these calculations is that there is no doubling of cells during the 8-hour pulsing window. Although that might be the case with some of the cells, many of the cells would have inevitably doubled and the observed total and labeled RNA could be coming from different number of cells from beginning of the pulsing to the end point. However, all these limitations apply to the estimations obtained by short-read TUC-seq and similar labeling techniques. While in this study we focus on labeling newly synthesized RNA using pulse labeling with 4SU, we could have instead performed a chase experiment to obtain degradation rates in situations where the main assumption would not hold.

Finally, the main advantage of using long-read sequencing for detection and quantification of recently transcribed genes is that it allows us to annotate the recently synthesized transcripts at isoform levels. Using this feature of long-read sequencing, we were able to identify a representative set of genes that, despite having rather ubiquitous expression across their isoforms, have substantially different transcription dynamics across isoforms. This could reflect the fact that some isoforms are required for a faster dynamic of a response whereas other isoforms are required to be more stable in order to confer robustness to some pathways. Having such resolution, one can infer the degradation rate, synthesis rate and the half-life of each of the isoforms and study the regulatory mechanism that affect these rates by integrating this data with other genomic assays such as miRNA-seq and ChIP-seq, and assays that focus on poly-A tails and 3’/5’-UTRs. In summary, Long-TUC-seq can robustly identify and quantify recently transcribed genes at the level of individual isoforms to shed light on differential isoform transcription and degradation rates.

## DISCUSSION

We would like to thank Melanie Oakes at UC Irvine Genomics High-Throughput Facility (GHTF) for her help with PacBio sequencing. This work was supported in part by grants from the National Institutes of Health (UM1HG009443 to A.M. and DP2GM119164 to RCS). RCS is a Pew Biomedical Scholar.

## SUPPLEMENTARY TABLES

**Table S1. Sequencing Statistics**

**Table S2. Gene level expression**

**Table S3. Transcript-level expression**

## REFERENCE

1. Munchel SE, Shultzaberger RK, Takizawa N, Weis K. Dynamic profiling of mRNA turnover reveals gene-specific and system-wide regulation of mRNA decay. Mol Biol Cell. 2011;22(15):2787–2795. doi: 10.1091/mbc.E11-01-0028

2. Lenstra TL, Rodriguez J, Chen H, Larson DR. Transcription Dynamics in Living Cells. Annu Rev Biophys. 2016;45(1):25–47. doi: 10.1146/annurev-biophys-062215-010838

3. Levine M, Tjian R. Transcription regulation and animal diversity. Nature. 2003;424(6945):147–151. doi: 10.1038/nature01763

4. Jiang S, Mortazavi A. Integrating ChIP-seq with other functional genomics data. Brief Funct Genomics. 2018;17(2):104–115. doi: 10.1093/bfgp/ely002

5. Buenrostro JD, Wu B, Chang HY, Greenleaf WJ. ATAC-seq: A method for assaying chromatin accessibility genome-wide. Curr Protoc Mol Biol. 2015;2015(January):21.29.1-21.29.9. doi: 10.1002/0471142727.mb2129s109

6. Maekawa S, Imamachi N, Irie T, et al. Analysis of RNA decay factor mediated RNA stability contributions on RNA abundance. BMC Genomics. 2015;16(1):1–19. doi: 10.1186/s12864-015-1358-y

7. Ghosh S, Jacobson A. mRNA decay modulates gene expression and controls its fidelity. Wiley Interdiscip Rev RNA. 2010;1(3):351–361. doi: 10.1007/springerreference_35999

8. Ule J, Jensen K, Mele A, Darnell RB. CLIP: A method for identifying protein-RNA interaction sites in living cells. Methods. 2005;37(4):376–386. doi: 10.1016/j.ymeth.2005.07.018

9. Alon S, Vigneault F, Eminaga S, et al. Barcoding bias in high-throughput multiplex sequencing of miRNA. Genome Res. 2011;21(9):1506–1511. doi: 10.1101/gr.121715.111

10. Core LJ, Waterfall JJ, Lis JT. Nascent RNA sequencing reveals widespread pausing and divergent initiation at human promoters. Science (80-). 2008;322(5909):1845–1848. doi: 10.1126/science.1162228

11. Mahat DB, Kwak H, Booth GT, et al. Base-pair-resolution genome-wide mapping of active RNA polymerases using precision nuclear run-on (PRO-seq). Nat Protoc. 2016;11(8):1455–1476. doi: 10.1038/nprot.2016.086

12. Nojima T, Gomes T, Grosso ARF, et al. Mammalian NET-seq reveals genome-wide nascent transcription coupled to RNA processing. Cell. 2015;161(3):526–540. doi: 10.1016/j.cell.2015.03.027

13. Churchman LS, Weissman JS. Native Elongating Transcript Sequencing ({NET}-seq). Curr Protoc Mol Biol. 2012;98(1):14.4.1--14.4.17. doi: 10.1002/0471142727.mb0414s98

14. Paulsen MT, Veloso A, Prasad J, et al. Coordinated regulation of synthesis and stability of RNA during the acute TNF-induced proinflammatory response. Proc Natl Acad Sci. 2013. doi: 10.1073/pnas.1219192110

15. Fuchs G, Voichek Y, Benjamin S, Gilad S, Amit I, Oren M. 4sUDRB-seq: measuring genomewide transcriptional elongation rates and initiation frequencies within cells. Genome Biol. 2014;15(5):R69. doi: 10.1186/gb-2014-15-5-r69

16. Schwalb B, Michel M, Zacher B, et al. TT-seq maps the human transient transcriptome. Science (80-). 2016;352(6290):1225–1228. doi: 10.1126/science.aad9841

17. Schofield JA, Duffy EE, Kiefer L, Sullivan MC, Simon MD. TimeLapse-seq: Adding a temporal dimension to RNA sequencing through nucleoside recoding. Nat Methods. 2018;15(3):221–225. doi: 10.1038/nmeth.4582

18. Herzog VA, Reichholf B, Neumann T, et al. Thiol-linked alkylation of {RNA} to assess expression dynamics. Nat Methods. 2017;14(12):1198–1204. doi: 10.1038/nmeth.4435

19. Gasser C, Delazer I, Neuner E, et al. Thioguanosine Conversion Enables mRNA-Lifetime Evaluation by RNA Sequencing Using Double Metabolic Labeling (TUC-seq DUAL). Angew Chemie - Int Ed. 2020. doi: 10.1002/anie.201916272

20. Russo J, Heck AM, Wilusz J, Wilusz CJ. Metabolic Labeling and Recovery of Nascent RNA to Accurately Quantify mRNA Stability. Methods. 2017;120:39–48. doi: 10.1016/j.physbeh.2017.03.040

21. Amarasinghe SL, Su S, Dong X, Zappia L, Ritchie ME, Gouil Q. Opportunities and challenges in long-read sequencing data analysis. Genome Biol. 2020;21(1):1–16. doi: 10.1186/s13059-020-1935-5

22. Wenger AM, Peluso P, Rowell WJ, et al. Accurate circular consensus long-read sequencing improves variant detection and assembly of a human genome Aaron. Nat Biotechnol. 2020;37(10):1155–1162. doi: 10.1038/s41587-019-0217-9.Accurate

23. Wyman D, Balderrama-Gutierrez G, Reese F, et al. A technology-agnostic long-read analysis pipeline for transcriptome discovery and quantification. bioRxiv. 2019:672931. doi: 10.1101/672931

24. Tardaguila M, De La Fuente L, Marti C, et al. Corrigendum: SQANTI: Extensive characterization of long-read transcript sequences for quality control in full-length transcriptome identification and quantification (Genome Research (2018) 28 (396-411) DOI: 10.1101/gr.222976.117). Genome Res. 2018;28(7):1096. doi: 10.1101/gr.239137.118

25. Tang AD, Soulette CM, Baren MJ Van, et al. Full-length transcript characterization of SF3B1 mutation in chronic lymphocytic leukemia reveals downregulation of retained introns. Nat Commun. 2020;11(2020):1–12. doi: 10.1038/s41467-020-15171-6

26. Wyman D, Mortazavi A. TranscriptClean: Variant-aware correction of indels, mismatches and splice junctions in long-read transcripts. Bioinformatics. 2019;35(2):340–342. doi: 10.1093/bioinformatics/bty483

27. Roberts A, Pachter L. Streaming fragment assignment for real-time analysis of sequencing experiments. Nat Methods. 2013;10(1):71–73. doi: 10.1038/nmeth.2251

